# Impaired Efflux of the Siderophore Enterobactin Induces Envelope Stress in *Escherichia coli*

**DOI:** 10.1101/779884

**Authors:** Randi L. Guest, Emily A. Court, Jayne L. Waldon, Kiersten A. Schock, Tracy L. Raivio

## Abstract

The Cpx response is one of several envelope stress responses that monitor and maintain the integrity of the gram-negative bacterial envelope. While several conditions that are known or predicted to generate misfolded inner membrane proteins activate the Cpx response, the molecular nature of the Cpx inducing cue is not yet known. Studies have demonstrated that mutation of multidrug efflux pumps activates the Cpx response in many gram-negative bacteria. In *Vibrio cholerae*, pathway activation is due to accumulation of the catechol siderophore vibriobactin. However, the mechanism by which the Cpx response is activated by mutation of efflux pumps in *Escherichia coli* remains unknown. Here we show that inhibition of efflux by deletion of *tolC*, the outer membrane channel of several multidrug efflux pumps, activates the Cpx response in *E. coli* as a result of impaired efflux of the siderophore enterobactin. Enterobactin accumulation in the *tolC* mutant reduces activity of the NADH oxidation arm of the aerobic respiratory chain. However, NADH dehydrogenase I, NADH dehydrogenase II, and cytochrome *bo*_*3*_ do not contribute to Cpx pathway activation in the *E. coli tolC* mutant. We show that the Cpx response down-regulates transcription of the enterobactin biosynthesis operon. These results suggest that the Cpx response promotes adaptation to envelope stress in enteric bacteria that are exposed to iron-limited environments, which are rich in envelope-damaging compounds and conditions.

## Introduction

In order for antimicrobial compounds to gain access to their cellular target, they must first cross one or more layers of the bacterial envelope. For gram-negative bacteria, this envelope consists of the outer membrane, the inner membrane, and the peptidoglycan sacculus located within the intervening periplasmic space (Silhavy et al., 2010). Antimicrobials that have crossed the envelope may be transported out of the cell via multidrug efflux pumps. Some multidrug efflux pumps interact with an outer membrane channel and periplasmic membrane fusion protein to form a tripartite protein complex that can directly transport toxic molecules from the cytoplasm or periplasm to the external environment, while others function as single component pumps that transport compounds from the cytoplasm to the periplasm (Li et al., 2015). Compounds transported to the periplasm via singlet efflux pumps may then move out of the cell through a tripartite machine (Lee et al., 2000; Tal and Schuldiner, 2009).

*Escherichia coli* encode several tripartite multidrug efflux systems, many of which use the same outer membrane channel, TolC (Li et al., 2015). Decades of research have shown that TolC is required for the efflux of a wide variety of dyes, detergents, and antibiotics. However, there is a growing body of evidence to suggest that TolC is also required for the secretion of endogenously produced metabolites. Intra- and extracellular concentrations of cysteine, indole, porphyrins, and siderophores are affected by loss of TolC or TolC-dependent efflux pumps (Bleuel et al., 2005; Horiyama and Nishino, 2014; Tatsumi and Wachi, 2008; Wiriyathanawudhiwong et al., 2009). Furthermore, accumulation of several metabolites increases expression of the TolC-dependent AcrAB multidrug efflux system as a compensatory mechanism to increase metabolite secretion (Helling et al., 2002; Ruiz and Levy, 2014). Blocking metabolite secretion by mutating *tolC* or TolC-dependent efflux systems increases sensitivity to cysteine, the siderophore enterobactin, and intermediates of heme biosynthesis, suggesting that metabolite accumulation is toxic (Tatsumi and Wachi, 2008; Vega and Young, 2014; Wiriyathanawudhiwong et al., 2009). In support of this hypothesis, numerous cellular stress responses are activated in bacteria lacking *tolC* (Guest and Raivio, 2016a; Rosner and Martin, 2009), including the Cpx envelope stress response.

Current evidence suggests that the Cpx envelope stress response functions to monitor and maintain the biogenesis of inner membrane proteins and protein complexes (Guest et al., 2017; Raivio, 2014; Vogt and Raivio, 2012). This response is controlled by a typical two-component signal transduction system consisting of the inner membrane bound sensor CpxA and the cytoplasmic response regulator CpxR (Dong et al., 1993; Weber and Silverman, 1988). In the presence of an inducing signal, CpxA autophosphorylates and the phosphate is then transferred to CpxR (Raivio and Silhavy, 1997). Once phosphorylated, CpxR functions as a transcription factor to activate the expression of genes associated with protein biogenesis and inner membrane integrity (Danese and Silhavy, 1997; 1998; Danese et al., 1995; Pogliano et al., 1997; Price and Raivio, 2009; Raivio et al., 2000; 2013), and repress the expression of genes that encode macromolecular envelope-localized protein complexes (Acosta et al., 2015; Dorel et al., 1999; Guest et al., 2017; Hernday et al., 2004; MacRitchie et al., 2008; McEwen and Silverman, 1980; Vogt et al., 2010). Once homeostasis is achieved, CpxA functions as a phosphatase to dephosphorylate CpxR and attenuate the response (Raivio and Silhavy, 1997).

Inhibition of efflux activates the Cpx response in several gram-negative bacteria, including *E. coli*, *Vibrio cholerae*, *Sinorhizobium meliloti*, and *Haemophilus ducreyi* (Acosta et al., 2014; Rinker et al., 2011; Rosner and Martin, 2013; Santos et al., 2010; Slamti and Waldor, 2009; Taylor et al., 2014), and is the most conserved Cpx inducing cue identified to date. Clues as to how impaired efflux activates the Cpx response have come from studies in *V. cholerae*. Activation of the Cpx pathway in *V. cholerae* lacking the TolC-dependent efflux system VexGH is suppressed when *V. cholerae* are grown in the presence of iron, suggesting that the metabolite responsible for activation of the *V. cholerae* Cpx response is produced when iron is limiting (Acosta et al., 2014). In a subsequent study, this metabolite was identified as the catechol siderophore vibriobactin (Kunkle et al., 2017). This study also found that the *V. cholerae* Cpx response is no longer activated in an efflux mutant when bacteria are grown anaerobically or when succinate dehydrogenase of the electron transport chain is disrupted. As such, it has been proposed that accumulation of vibriobactin activates the Cpx response via the electron transport chain.

It is thought that vibriobactin production is limited to a small number of *Vibrio* species (Thode et al., 2018; Wyckoff et al., 2001). As such, the mechanism by which inhibition of efflux activates the Cpx response in *E. coli* remains to be determined. In this study, we show that the catechol siderophore enterobactin is required for activation of the Cpx response in *E. coli* lacking *tolC*, suggesting that envelope damage inflicted by impaired secretion of siderophores is a conserved Cpx inducing signal. While enterobactin was found to decrease activity of the NADH oxidation arm of the aerobic electron transport chain in the *tolC* mutant, neither NADH dehydrogenase I, NADH dehydrogenase II, nor cytochrome *bo*_*3*_ contribute to activation of the Cpx response in this background. Finally, we provide evidence to suggest that activation of the Cpx response facilitates adaptation to toxic envelope stresses, such as enterobactin accumulation, by down-regulating the transcription of genes involved in enterobactin biosynthesis.

## Results

### Iron limitation induces the Cpx response in the *tolC* mutant

To confirm that inhibition of efflux activates the Cpx response in *E. coli*, we measured Cpx pathway activity in a *tolC* mutant using a *cpxP-lacZ* transcriptional reporter. No change in *cpxP-lacZ* reporter activity was observed when *E. coli* were grown in LB (figure 1A). This was surprising given that under similar growth conditions, expression of the periplasmic chaperone Spy has been shown to increase in a *tolC* mutant via the Cpx response (Acosta et al., 2014; Rosner and Martin, 2013). Nonetheless, when *E. coli* were grown in M9 minimal medium we observed a nearly eleven-fold increase in *cpxP-lacZ* activity in the *tolC* mutant (figure 1A). This increase was abolished in *E. coli* lacking *cpxA* (figure 1B), suggesting that inhibition of efflux generates envelope stress that is sensed by CpxA. These results suggest that the metabolite(s) responsible for activating the Cpx response in the *E. coli tolC* mutant is produced in minimal medium, but not in rich medium.

**Figure 1:**
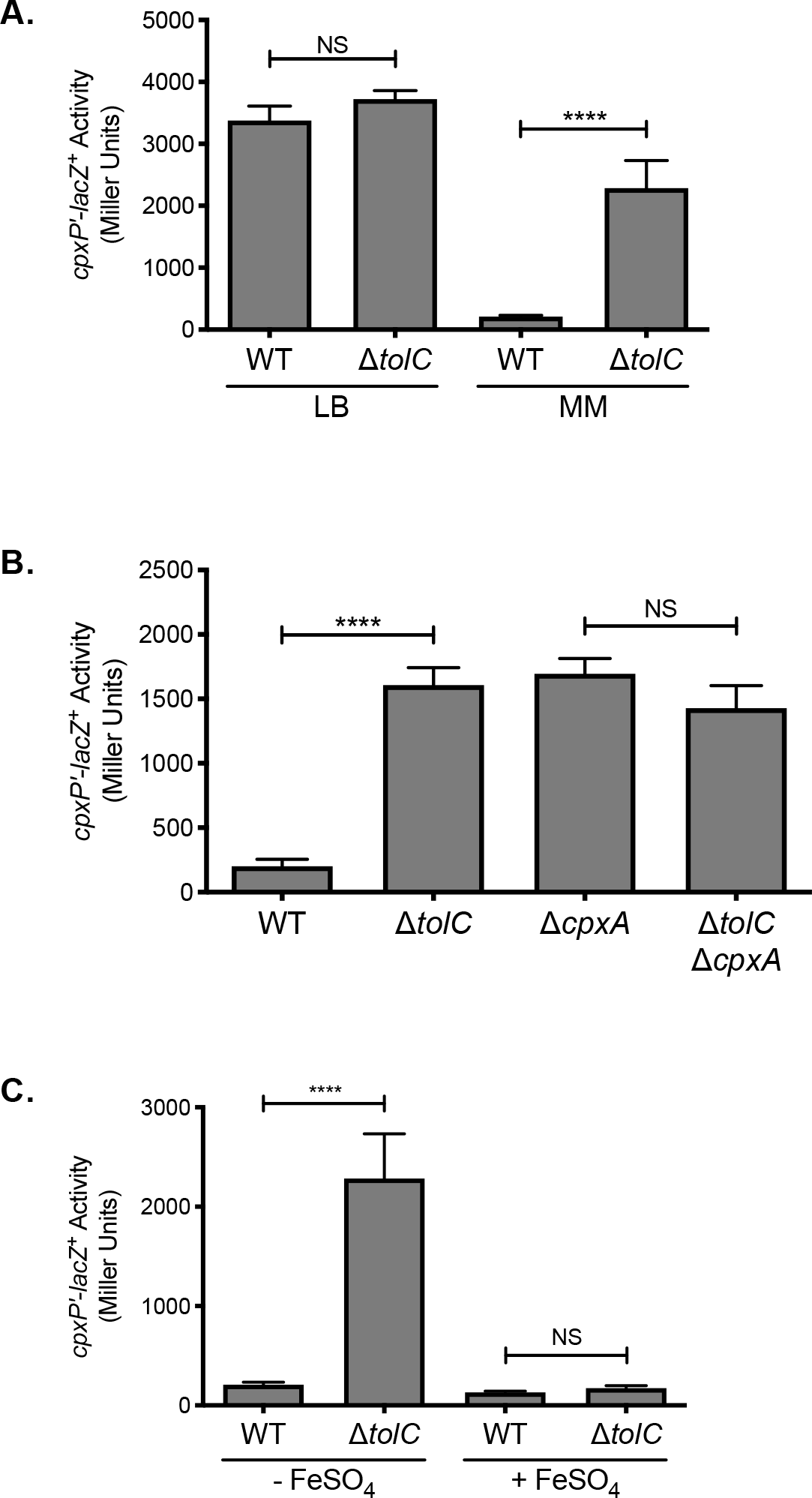
Deletion of *tolC* activates the *E. coli* Cpx response under iron-deplete conditions. (A) Wildtype and *tolC* mutant *E. coli* MC4100 strains carrying the chromosomal *cpxP-lacZ* transcriptional reporter were subcultured into Lennox broth (LB) or M9 minimal medium (MM) after overnight growth in LB medium and grown for twenty hours at 37°C. (B) *cpxP-lacZ* activity in wildtype *E. coli* strain MC4100, the *tolC* and *cpxA* single mutants, and the *tolC cpxA* double mutant, grown in M9 minimal medium. (C) Expression of the *cpxP-lacZ* in wildtype and *tolC* mutant *E. coli* MC4100 strains subcultured into M9 minimal medium with (+) or without (−) 80μM FeSO_4_ from overnight cultures grown in LB medium. Bacteria were grown for 20 hours at 37°C. To measure *cpxP-lacZ* expression, cells were lysed with chloroform and SDS, and β-galactosidase levels were measured after addition of ONPG in a 96 well plate as described in the materials and methods. Data represent the means and standard deviations of three biological replicates. Asterisks indicate a statistically significant difference from the indicated wildtype control (****, *P* ≤ 0.0001 [one-way ANOVA with Sidak’s post-hoc test]). NS indicates no statistically significant difference in *cpxP-lacZ* reporter activity. WT, wildtype.

Previous results from our lab have shown that impaired efflux activates the *V. cholerae* Cpx response when iron is limiting (Acosta et al., 2014). As such, we hypothesized that iron may be involved in activation of the Cpx response in the *E. coli tolC* mutant. As observed previously, mutation of *tolC* resulted in approximately an eleven-fold increase in *cpxP-lacZ* activity when *E. coli* were grown in the iron deplete M9 minimal medium (figure 1C). However, when the *tolC* mutant was grown in M9 minimal medium supplemented with 80μM FeSO_4_, activation of the Cpx response was no longer observed (figure 1C). These results suggest that the metabolite(s) responsible for activating the Cpx response in *E. coli* lacking *tolC* is produced during iron deprivation.

### Enterobactin activates the Cpx response in the E. coli *tolC* mutant

Several lines of evidence implicate the siderophore enterobactin in activation of the *E. coli* Cpx response in the *tolC* mutant. First, enterobactin is synthesized in response to iron starvation and its production is repressed in the presence of iron by the master iron regulator Fur (Bagg and Neilands, 1985; Brickman et al., 1990). Second, enterobactin is secreted into the environment via TolC (Bleuel et al., 2005) and in the absence of TolC, enterobactin accumulates in the periplasm (Vega and Young, 2014). Finally, enterobactin is structurally similar to vibriobactin (Griffiths et al., 1984), the metabolite responsible for activating the Cpx response in *V. cholerae* efflux mutants (Kunkle et al., 2017). To test whether accumulation of enterobactin is responsible for activating the Cpx response in absence of *tolC* in *E. coli*, we disrupted enterobactin biosynthesis genetically by deleting *entC* (Young et al., 1971). Unlike the *tolC* single mutant where *cpxP-lacZ* activity was increased eleven-fold, there was no increase in *cpxP-lacZ* activity in the *tolC entC* double mutant (figure 2A). Furthermore, addition of exogenous enterobactin to the medium restored Cpx pathway activation in the *tolC entC* double mutant (figure 2B). Together, these results suggest that enterobactin is responsible for activating the Cpx stress response in the *E. coli tolC* mutant and demonstrate that accumulation of catechol siderophores generates a Cpx inducing signal that is conserved in *V. cholerae* and *E. coli*.

**Figure 2:**
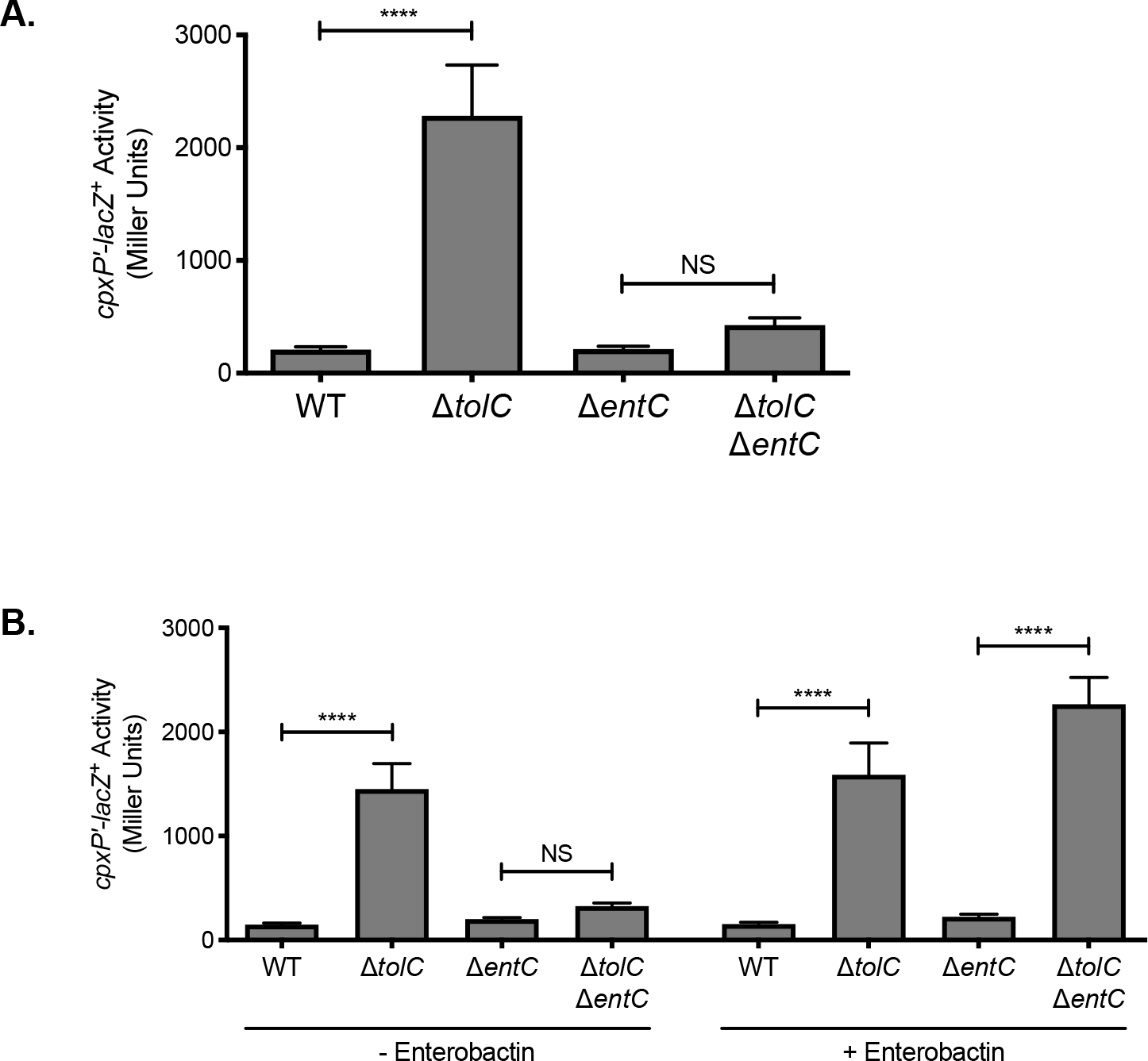
Enterobactin activates the Cpx response in the *tolC* mutant. (A and B) Expression of the chromosomal *cpxP-lacZ* transcriptional reporter in wildtype *E. coli* strain MC4100, the *tolC* and *entC* single mutants, and the *tolC entC* double mutant. Strains were grown overnight in LB and then subcultured into (A) M9 minimal medium, or (B) M9 minimal medium with (+) or without (−) 10μM enterobactin. As enterobactin is dissolved in 42% DMSO, an equivalent volume of 42% DMSO was added to cultures without (−) enterobactin. Bacteria were grown for 20 hours at 37°C. Cells were lysed with chloroform and SDS, and β-galactosidase levels were measured after addition of ONPG in a 96 well plate as described in the materials and methods section. Data show means and standard deviations of three biological replicates. Asterisks indicate a statistically significant difference from the indicated Wildtype control (****, *P* ≤ 0.0001 [one-way ANOVA with Sidak’s post-hoc test]). NS indicates no statistically significant difference in *cpxP* reporter activity. WT, wildtype.

### Impaired secretion of enterobactin decreases NADH oxidase activity

We next sought to determine whether other phenotypes associated with the *tolC* mutant are due to impaired secretion of enterobactin. While *tolC* is not essential for growth in rich medium, the growth rate of *tolC* deficient *E. coli* is substantially reduced in minimal medium (Dhamdhere and Zgurskaya, 2010; Vega and Young, 2014). One study attributed this phenotype to reduced activity of NADH dehydrogenase of the electron transport chain (Dhamdhere and Zgurskaya, 2010), while another found that the growth defect could be suppressed by the addition of iron (Vega and Young, 2014). Together, these results suggest that enterobactin may reduce NADH dehydrogenase activity in the *tolC* mutant. To examine this possibility, we measured NADH oxidase activity in the *tolC* and *entC* single mutants and *tolC entC* double mutant by measuring the rate of oxygen consumption using β-NADH as the electron donor. As expected, oxygen consumption is reduced in the *tolC* mutant compared to the wildtype (figure 3). However, oxygen consumption in the *tolC entC* double mutant is similar to that of the WT and *entC* single mutant (figure 3). These results are consistent with the hypothesis that enterobactin is responsible for reduced NADH dehydrogenase activity in the *tolC* mutant.

**Figure 3:**
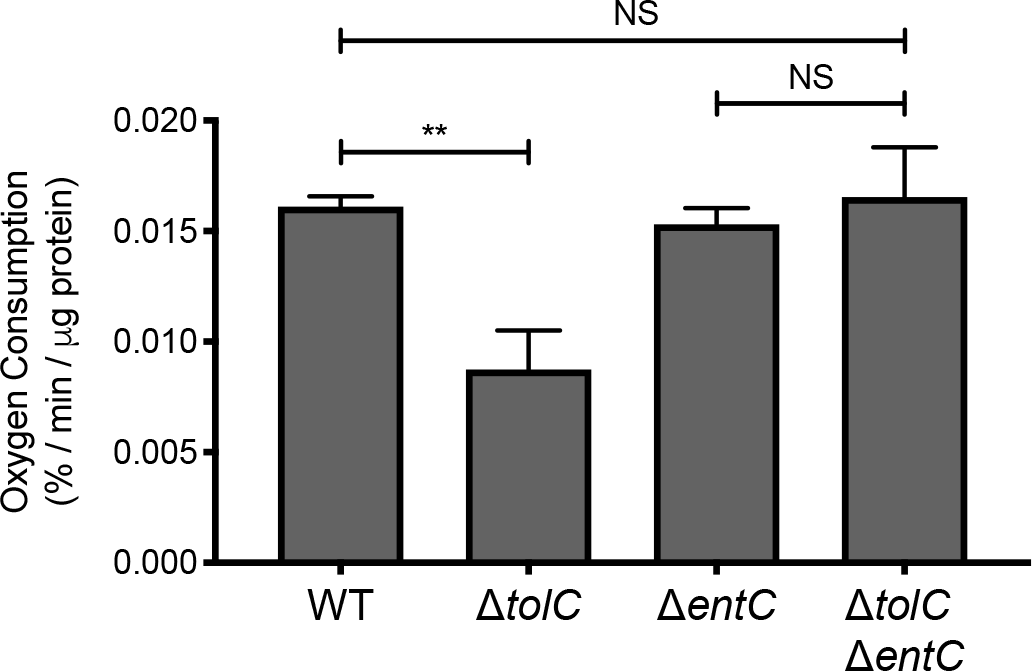
Enterobactin reduces NADH oxidase activity in the *tolC* mutant. Wildtype (WT) *E. coli* strain MC4100, the *tolC* and *entC* single mutants, and the *tolC entC* double mutant were subcultured into M9 minimal medium containing 0.4% glucose after overnight growth in LB medium and grown for 20 hours at 37°C. Bacteria were collected by centrifugation and washed once in 50mM MES buffer, pH 6.0. Bacteria were then pelleted by centrifugation, weighed, and resuspended in 1mL of 50mM MES buffer, pH 6.0. Bacteria were lysed by sonication. 100μL of cell lysate was diluted in 890μL 50mM MES buffer pre-warmed to 30°C in a 1mL microrespiration chamber. Diluted lysate was covered in light mineral oil to prevent oxygen from dissolving into the system. Oxygen concentration was measured every 30 seconds for 10-15 minutes after the addition of 100μM β-NADH at 30°C using oxygen MicroOptode sensor (Unisense). Oxygen concentration at each time point was standardized to the concentration present just prior to the addition of β-NADH. The rate of oxygen concentration per μg of total protein was calculated as described in the materials and methods section. Data represent the means and standard deviations of three biological replicates. Asterisks indicate a statistically significant difference from the indicated strain (**, *P* ≤ 0.01 [one-way ANOVA with Sidak’s post-hoc test]). NS indicates no statistically significant difference in the rate of oxygen consumption.

### The Cpx response in the *tolC* mutant remains active in the absence of aerobic electron transport chain components

*E. coli* encode two NADH dehydrogenase isoenzymes that can oxidize β-NADH, NADH dehydrogenase I (NDH-I) and NADH dehydrogenase II (NDH-II) (Matsushita et al., 1987). Under aerobic conditions, electrons released from β-NADH by NDH-I or NDH-II are first transferred to ubiquinone and then to a terminal oxidase, such as cytochrome *bo*_3_. Given that activity of the NADH oxidation arm of the aerobic respiratory chain is impaired in the *tolC* mutant (figure 3), and that NDH-I and cytochrome *bo*_*3*_ contribute to Cpx pathway activity in enteropathogenic *E. coli* (Guest et al., 2017), we next asked whether NDH-I, NDH-II, or cytochrome *bo*_*3*_ are required for activation of the Cpx response in the *tolC* mutant. To test this hypothesis, we mutated each of these electron transport chain components in *E. coli* containing or lacking *tolC* and measured Cpx pathway activity using the *cpxP-lacZ* transcriptional reporter. As many of the mutants grew poorly in minimal glucose broth, we determined *cpxP-lacZ* expression from bacteria grown on M9 minimal agar medium. Deletion of *tolC* activated the Cpx response under these conditions, evidenced by a 5.5-fold increase in *cpxP-lacZ* activity in the *tolC* mutant compared to the wildtype (figure 4). Furthermore, *cpxP-lacZ* expression was increased to a lesser extent in the *tolC entC* double mutant than in the *tolC* single mutant (figure S1), confirming that activation of the Cpx response in the *tolC* mutant is dependent on enterobactin under these conditions as well. The Cpx response was activated to a similar extent in the *tolC nuoA-N* (NADH dehydrogenase I), *tolC ndh* (NADH dehydrogenase II), and *tolC cyoA-E* (cytochrome *bo*_*3*_), double mutants as in the *tolC* single mutant (figure 4), suggesting that these components are not required for activation of the Cpx response in *E. coli* lacking *tolC.*

**Figure 4:**
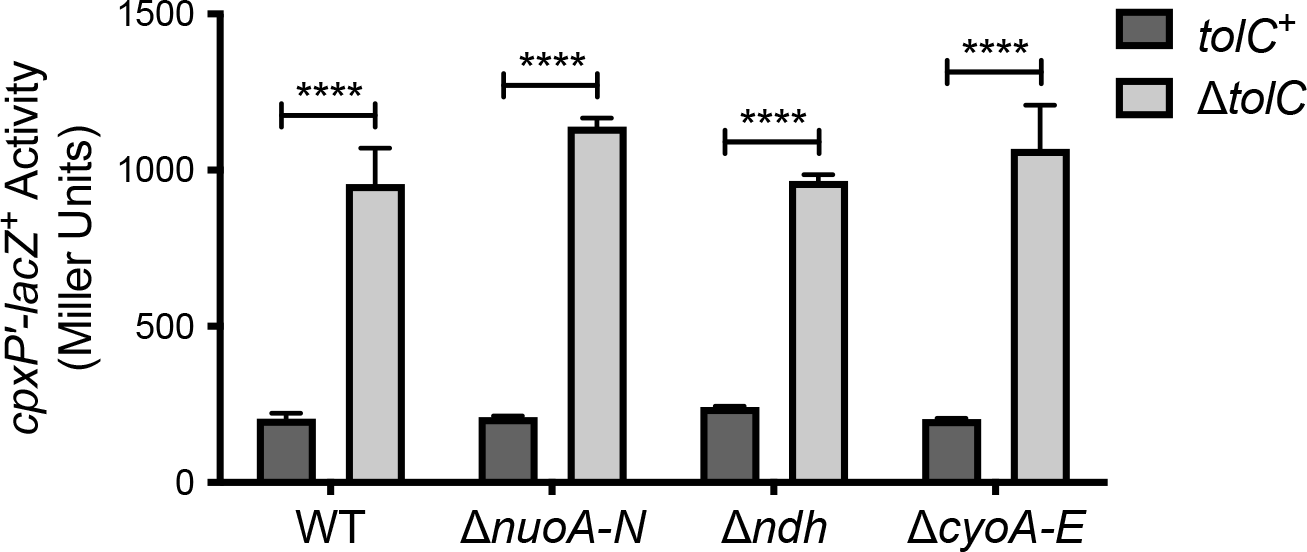
Activation of the Cpx response by deletion of *tolC* does not require NDH-I, NDH-II, or cytochrome *bo*_*3*_. After overnight growth in LB medium, bacteria containing the *cpxP-lacZ* transcriptional reporter were washed once, and resuspended in, phosphate-buffered saline. 10μL of culture was spotted onto M9 minimal medium agar containing 0.4% glucose and grown at 37°C for 24 hours. Bacteria were collected using plastic inoculating loops and resuspended in 1 × Z buffer. *cpxP-lacZ* activity was then measured as described in the materials and methods section. The strains shown are TR50 and the isogenic Δ*tolC,* Δ*nuoABCDEFGHIJKLMN∷kan*, Δ*ndh∷kan,* and Δ*cyoABCDE∷kan* mutants, and the Δ*tolC* Δ*nuoABCDEFGHIJKLMN∷kan,* Δ*tolC* Δ*ndh∷kan*, and Δ*tolC* Δ*cyoABCDE∷kan* double mutants. Data represent the means and standard deviations of three biological replicates. Asterisks indicate a statistically significant difference from the control strain containing a wildtype copy of *tolC* (*tolC^+^*) (****, *P* ≤ 0.0001 [two-way ANOVA with Sidak’s post-hoc test]).

### Regulation of enterobactin biosynthesis genes by the Cpx response

Previous microarray experiments performed to identify members of the Cpx regulon in enteropathogenic *E. coli* (EPEC) and *E. coli* K-12 strain MC4100 found that expression of several genes involved in enterobactin biosynthesis are decreased upon activation of the Cpx response, including *entA, entB, entC, and entE* (Raivio et al., 2013). To confirm regulation of the enterobactin biosynthesis genes by the Cpx response, we constructed luminescent transcriptional reporters of EPEC and MC4100 *entCEBA* expression. Activity of each lux reporter was analyzed in wildtype EPEC or MC4100, mutants containing the *cpxA24* allele that constitutively activates the Cpx response, and in *E. coli* lacking the Cpx response. Mutational activation of the Cpx response in EPEC resulted in a 4.0-fold decrease in activity of the EPEC *entCEBA-lux* reporter (figure 5A). No change in reporter activity was observed in the EPEC Δ*cpxRA* mutant (figure 5A), suggesting that basal expression of the enterobactin biosynthesis genes is not affected by loss of the Cpx response. Likewise, activation of the Cpx response in MC4100 led to a 6.2-fold decrease in activity of the MC4100 *entCEBA-lux* reporter in comparison to the wildtype (figure 5B). No change in MC4100 *entCEBA-lux* activity was observed in the MC4100 *cpxR* mutant (figure 5B). These data suggest that the transcription of genes involved in enterobactin biosynthesis is repressed upon activation of the Cpx response in EPEC and MC4100.

**Figure 5:**
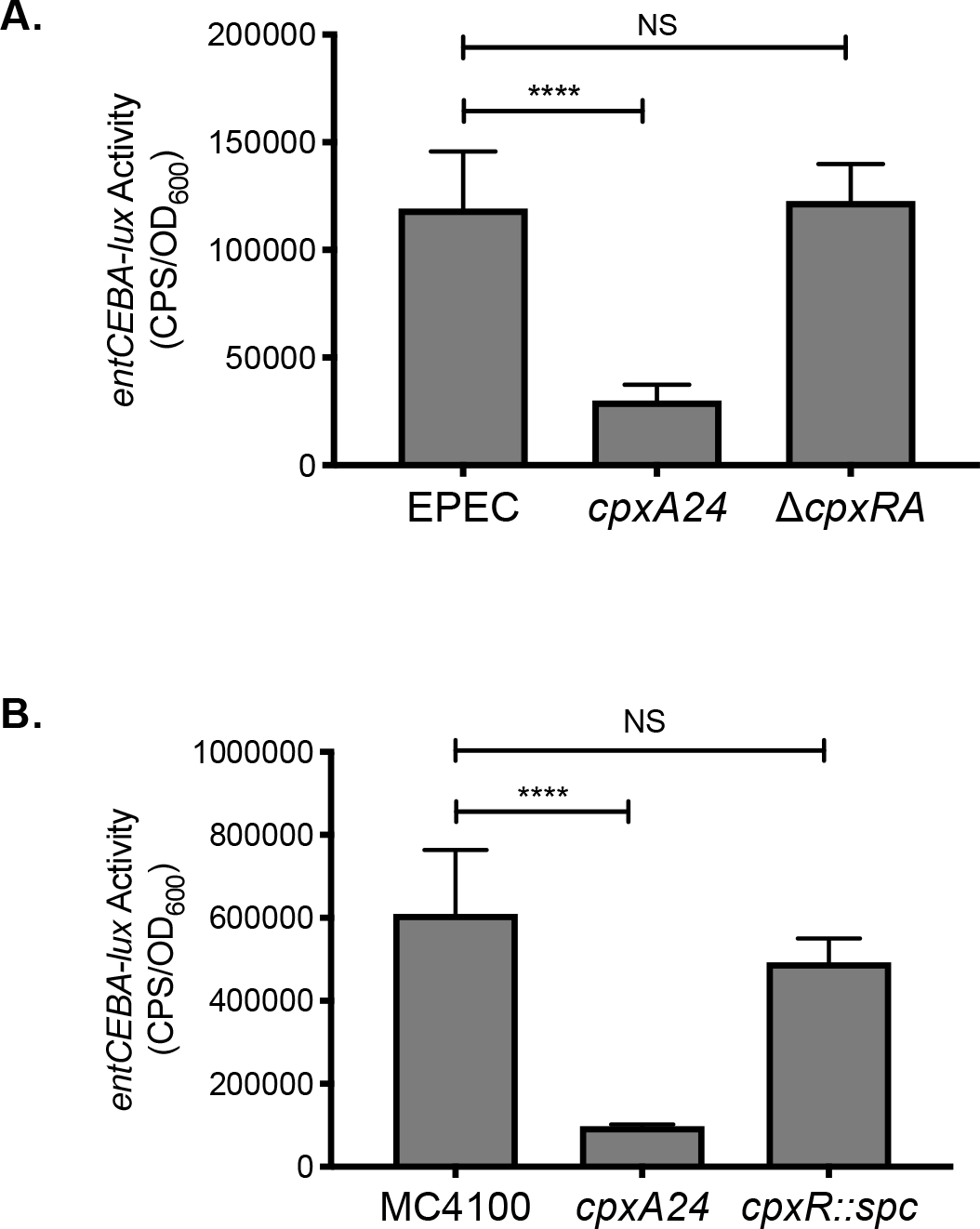
Transcription of the *entCEBA* operon is repressed by the Cpx response. Bacteria were grown overnight in LB broth at 30°C with shaking. The following day, bacteria were subcultured into M9 minimal medium supplemented with 0.4% glucose, 5.34mM isoleucine, and 6.53mM valine and grown for 8 hours at 30°C with shaking. 200μL of culture was transferred to a black-wall 96 well plate and luminescence (expressed in counts per second [CPS]) and OD_600_ were read, and *entCEBA-lux* activity was calculated as described in the materials and methods section. (A) Activity of the EPEC *entCEBA-lux* reporter in wildtype EPEC and EPEC containing the *cpxA24* or Δ*cpxRA* mutation. (B) Activity of the MC4100 *entCEBA-lux* reporter in wildtype MC4100 and MC4100 containing the *cpxA24* or *cpxR∷spc* mutation. Asterisks indicate a statistically significant difference between the indicated strains (****, *P* ≤ 0.0001 [one-way ANOVA with Sidak’s post-hoc test]). NS indicates no statistically significant difference in *entCEBA-lux* reporter activity.

Intriguingly, we observed differences in basal *entCEBA* expression between MC4100 and EPEC. Activity of the native *entCEBA-lux* reporter in MC4100 was 5.1-fold higher than activity of the native *entCEBA-lux* reporter in EPEC (figure S2A). The promoter regions of the *entCEBA* operon are substantially different between MC4100 and EPEC (figure S2B). In addition to several base pair substitutions, there is an 186bp deletion in the promoter region of the EPEC *entCEBA* operon. There are two possible explanations for the difference in the basal levels of *entCEBA* transcription in MC4100 and EPEC. The first possibility is that the difference in the DNA sequence of the EPEC *entCEBA* promoter decrease basal transcription of the *entCEBA* operon. If true, we would expect that activity of the EPEC *entCEBA-lux* reporter would decrease in MC4100 and that activity of the MC4100 *entCEBA-lux* reporter would increase in EPEC. The second possibility is that activity of transcription factors that regulate *entCEBA* transcription is different between EPEC and MC4100. Here, expression of the EPEC and MC4100 *entCEBA-lux* reporters would both be similar in EPEC. Likewise, expression of both reporters would be similar in MC4100. We found that activity of the EPEC and MC4100 *entCEBA-lux* reporters were similar in EPEC, suggesting that the difference in *entCEBA* transcription is not due to differences in the DNA sequence of the *entCEBA* promoter regions (figure S2A). Furthermore, we found that activity of the EPEC *entCEBA-lux* reporter in MC4100 is actually increased approximately two-fold in comparison to the activity of the MC4100 *entCEBA-lux* reporter in MC4100 (figure S2A). Accordingly, these results suggest that expression of the *entCEBA* operon is decreased in EPEC through changes in activity of transcriptional regulators.

## Discussion

Multidrug efflux pumps export a wide range of antimicrobial compounds and thus play a major role in resistance of gram-negative bacteria to various antibiotics. However, several studies have revealed that multidrug efflux pumps are involved in cellular processes beyond antibiotic resistance, including cell division, biofilm formation, pathogenesis, cell communication, oxidative and nitrosative stress resistance, and envelope biogenesis (Baranova, 2016; Fahmy et al., 2016; Poole and Fruci, 2016). It has been proposed that drug efflux pumps function to secrete toxic endogenous metabolites that disrupt cellular integrity (Helling et al., 2002; Rosner and Martin, 2009). In this study, we report that the catechol siderophore enterobactin is responsible for activating the Cpx envelope stress response in *E. coli* lacking TolC, the outer membrane channel of several multidrug efflux systems (figure 6). While the mechanism by which impaired secretion of enterobactin activates the Cpx response remains to be determined, our data suggest that NDH-I, NDH-II, and cytochrome *bo*_*3*_ are not involved.

**Figure 6.**
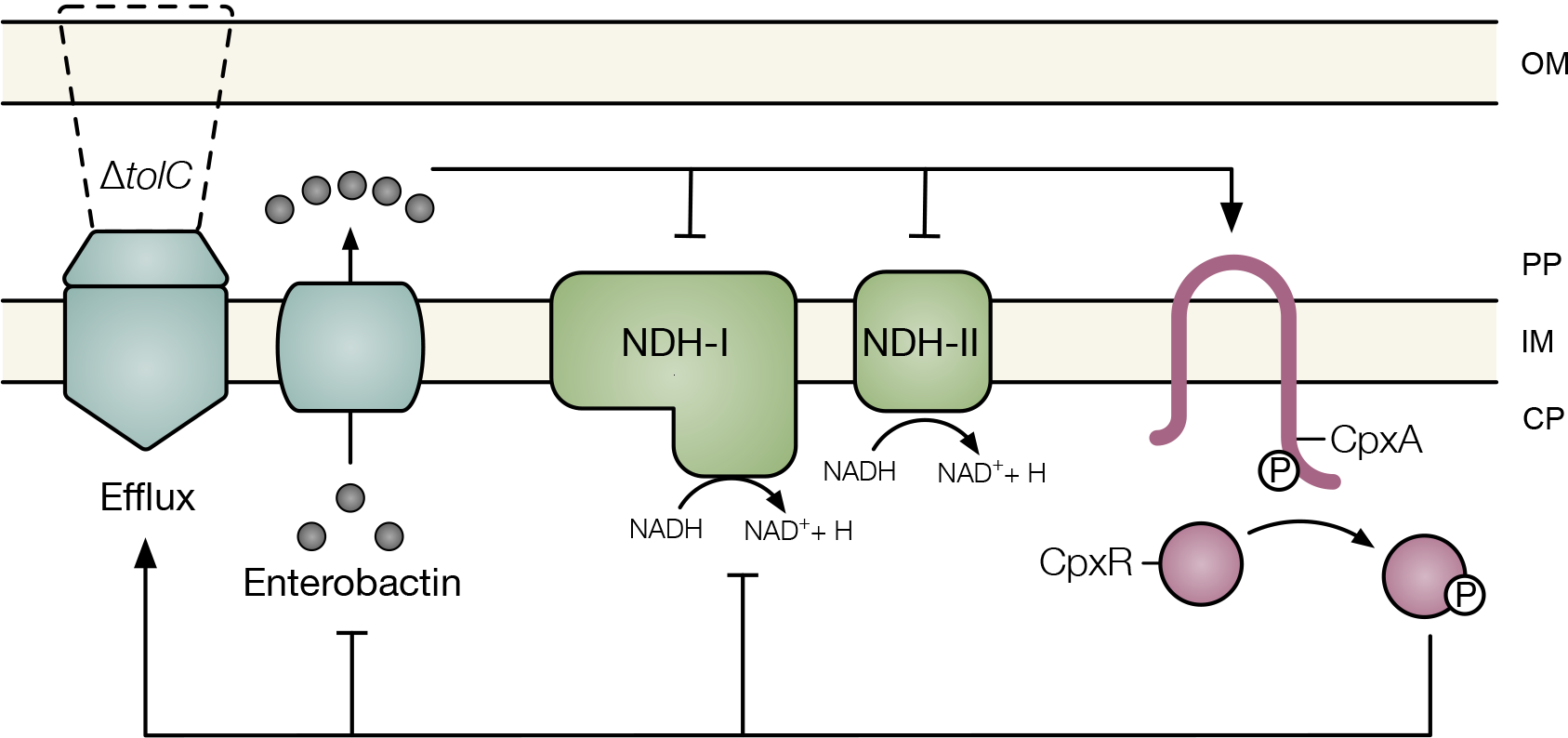
Model of the association between the impaired efflux and the Cpx envelope stress response. Under conditions in which efflux through TolC is compromised, the siderophore enterobactin accumulates within the periplasm of *E. coli*. Impaired secretion of enterobactin activates the Cpx response and disrupts NADH oxidase activity. Cpx regulation of multidrug efflux pumps, respiratory complexes, and enterobactin biosynthesis may provide an adaptive response to enterobactin accumulation. OM, outer membrane; PP, periplasm; IM, inner membrane; CP, cytoplasm; Q, quionone; P, phosphate.

The enterobactin secretion pathway begins in the cytoplasm, where enterobactin is synthesized (reviewed in Raymond et al., 2003). Cytoplasmic enterobactin is then transported to the periplasm via the singlet efflux pump EntS (Furrer et al., 2002). Once in the periplasm, enterobactin is secreted into the environment by one of several TolC-dependent tripartite efflux systems (Bleuel et al., 2005). Enterobactin that has bound to iron is brought back into the cell through the TonB-dependent outer membrane channel FepA, translocated across the periplasm by FepB, and moved into the cytoplasm by the FepCDG inner membrane transporter (reviewed in (Guerinot, 1994). Next, the iron-enterobactin complex is hydrolyzed by Fes, which liberates the bound iron and breaks down enterobactin into mono-di- and/or tri-dihydroxybenzoylserine (Brickman and McIntosh, 1992; O’Brien et al., 1971). While our data suggest that impaired secretion of enterobactin induces the Cpx response, the point in the impaired secretion pathway at which envelope stress is generated is unknown. As deletion of *tolC* does not activate the Cpx pathway in *E. coli* lacking CpxA, the stress that activates the Cpx response is likely located in the periplasm or inner membrane. In this regard, the toxic defects associated with the *E. coli tolC* mutant grown under iron-limiting conditions are due to the accumulation of periplasmic enterobactin (Vega and Young, 2014). Notably, basal levels of enterobactin in *E. coli* containing a functional TolC protein do not affect activity of the Cpx response, as *cpxP-lacZ* expression was unchanged in the *entC* single mutant compared to wildtype (figure 2A). As such, we hypothesize that it is the aberrant accumulation of periplasmic enterobactin in *E. coli* lacking *tolC* that activates the Cpx response.

Whether impaired secretion of enterobactin is responsible for the pleiotropic phenotypes displayed by bacteria lacking *tolC* remains to be determined. At present, it is known that enterobactin induces the growth and morphological defects associated with *tolC* mutants grown in iron-limited medium, but is not required for antibiotic hypersensitivity (Vega and Young, 2014). In a previous study, deletion of *tolC* in *E. coli* was found to decrease NADH oxidase activity when cells were grown in minimal medium, but not rich medium (Dhamdhere and Zgurskaya, 2010). Here, we show that this is due to impaired secretion of enterobactin. There are two possible explanations as to why enterobactin may reduce NADH oxidase activity in the *tolC* mutant. The first possibility is that enterobactin damages one or both of the NADH dehydrogenase protein complexes, thus reducing their ability to oxidize NADH. The second possibility is that activation of the Cpx response, or another regulatory system, by deleting *tolC* reduces expression of NDH-I or NDH-II. In support of this hypothesis, CpxR has been shown to directly repress the transcription of the operon encoding NDH-I (Guest et al., 2017). At this point, we are unable to distinguish between these two possibilities.

Although we and others have determined that siderophore accumulation is responsible for activating the Cpx response in the *tolC* mutant, the nature of the Cpx-inducing stress that is generated under this condition is unknown. Given that activation of the Cpx response in the *V. cholerae tolC* mutant can also be prevented by growing cells anaerobically or by disrupting succinate dehydrogenase, it has been proposed that siderophores chelate iron from the iron-containing cofactors present in aerobic respiratory complexes (Kunkle et al., 2017). Damaged respiratory complexes may either activate the Cpx response directly or increase formation of reactive oxygen species that then generate a Cpx-inducing signal (Kunkle et al., 2017). Here, we found that NDH-I, NDH-II, and cytochrome *bo*_*3*_ are not required for activation of the Cpx response in the *E. coli tolC* mutant. We were unable to accurately assess the role of succinate dehydrogenase in the activation of the Cpx response in the *E. coli* lacking *tolC*, as the succinate dehydrogenase mutant grew very poorly in iron-limiting minimal medium (data not shown). Together, these results suggest that succinate dehydrogenase, but not other components of the aerobic electron transport chain, contributes to activation of the Cpx response in bacteria lacking *tolC*. Whether and how siderophores impair succinate dehydrogenase remains to be determined.

In agreement with previous transcriptomic data, we found that the Cpx stress response represses expression of the genes for enterobactin biosynthesis in EPEC and *E. coli* K-12. We also found that basal transcription of the *entCEBA* operon is decreased in EPEC in comparison to MC4100, likely due to changes in activity of transcription factors in EPEC. As expression of the enterobactin biosynthesis genes is regulated in response to intracellular iron concentrations (Bagg and Neilands, 1985; Brickman et al., 1990), it is possible that intracellular iron concentrations are different in EPEC and MC4100. Alternatively, it is possible that pathogens such as EPEC more tightly control regulation of iron metabolism to facilitate host colonization.

In addition to regulation of enterobactin biosynthesis, other processes regulated by the Cpx response could facilitate adaptation to the stress caused by enterobactin accumulation. Activation of the Cpx response increases expression of multidrug efflux pumps in *E. coli*, *V. cholerae*, *Pseudomonas aeruginosa*, and *Klebsilla pneumoniae* (Guest and Raivio, 2016b; Tian et al., 2016), which could lead to increased efflux of periplasmic enterobactin. Indeed, the Cpx-regulated expression of the VexGH efflux pump, which is required for vibriobactin secretion, supports this hypothesis (Kunkle et al., 2017). Furthermore, the Cpx response in *E. coli* represses expression of several components of the electron transport chain, including succinate dehydrogenase (Guest et al., 2017; Raivio et al., 2013). As such, activation of the Cpx response would decrease expression of the potential target of enterobactin-mediated stress. Through decreased enterobactin biogenesis, increased efflux, and decreased expression of the target respiratory complex, activation of the Cpx response could mount an effective adaptive response to the stress exerted by enterobactin accumulation (figure 6).

Several noxious compounds secreted by TolC are present in the host environment. Enteric bacteria such as *E. coli* and *V. cholerae* are exposed to host-produced factors such as bile and cationic antimicrobial peptides, as well as antibiotics produced by competing members of the intestinal microbiome. Furthermore, *E. coli* and *V. cholerae* likely synthesize and secrete siderophores in response to the iron-poor environment within the host. As a large number of noxious compounds present *in vivo* require TolC for secretion, it is possible that they accumulate within the cell faster than can be effluxed through TolC. Activation of the Cpx response could provide protection against the surges in periplasmic siderophore concentrations that occur under these conditions.

## Experimental Procedures

### Bacterial strains and growth conditions

All bacterial strains and plasmids used in the course of this study are listed in table S1. Bacteria were grown in either lennox broth (LB, 10g/L bactotryptone [Difco], 5g/L yeast extract [Difco], and 5g/L NaCl) or M9 minimal medium (Difco) containing 0.4% glucose at 37°C with shaking at 225 rpm. Bacteria were grown at 30°C for experiments that included strain TR10 or ALN195. Antibiotics were added as necessary to the following concentrations: amikacin (Amk), 3μg mL^−1^; ampicillin (Amp), 100μg mL^−1^; kanamycin (Kan), 50μg mL^−1^; spectinomycin (Spc), 25μg mL^−1^; streptomycin (Str), 50μg mL^−1^. All chemicals were purchased from Sigma-Aldrich unless otherwise stated.

### Strain and plasmid construction

Strains EC3, EC4, RG244, RG249, and RG250 were constructed by P1 transduction (Silhavy et al., 1984). Donor strains, in which the *tolC*, *entC, ndh,* or *cpxA* open-reading frame was replaced with the kanamycin resistance cassette, were obtained from the Keio library (Baba et al., 2006). The kanamycin resistance cassette in the *tolC* gene was removed by FLP/FRT mediated recombination to produce an in-frame, markerless deletion as described in Hoang *et al*. (1998). All mutations were confirmed by PCR.

Deletion of the *nuoABCDEFGHIJKLMN* and the *cyoABCDE* operons in DY378 was performed by lamba-red recombinase as previously described (Thomason et al., 2007). Primer sequences were obtained from (Baba et al., 2006) (table S2). The DNA sequence of the K12nuoKOF primer corresponds to the 5’ primer used to delete *nuoA* in (Baba et al., 2006), while sequence of the K12nuoKOR primer corresponds to the 3’ primer used to delete *nuoN* (Baba et al., 2006). Primer K12-cyoKOF corresponds to the 5’ primer used to delete *cyoA* in (Baba et al., 2006), and primer K12-cyoKOR corresponds to the 3’ primer used to delete *cyoE* (Baba et al., 2006). PCR was performed using high-fidelity Phusion DNA polymerase (ThermoFisher) according to the manufacturers specifications with the addition of 20% betaine. K12nuoKOF and K12nuoKOR, or K12-cyoKOF and K12-cyoKOR, were used to amplify the FRT-flanked kanamycin resistance cassette from the keio library (Baba et al., 2006). DNA was separated by electrophoresis on a 1% agarose gel. A DNA fragment approximately the size of the kanamycin resistance cassette was extracted and cleaned using the GeneJet gel purification kit (Fermentas). These DNA fragments were used to delete the *nuoA-N* or *cyoA-E* locus in *E. coli* strain DY378, which encodes the lambda-red recombinase system from a temperature sensitive promoter (Thomason et al., 2007; Yu et al., 2000). Briefly, DY378 was grown to an OD_600_ of 0.4-0.5 in 35mL of LB in a 250mL Erlenmeyer flask at 30°C with shaking at 225 rpm. Half of this culture was then transferred to a 125mL flask and incubated in a 42°C shaking water bath for 15 minutes, while the other half was incubated at 30°C as before. Cells were washed three times in sequentially lower volumes of ice-cold distilled water, terminating with cells resuspended in 200μL ice-cold distilled water. 100ng or 300ng of purified kanamycin resistance cassette DNA with homologous ends to the *nuo* or *cyo* operon were electroportated into DY378 and cells were recovered at 30°C with shaking for 2 hours. Recombinants were selected for on LB agar supplemented with kanamycin. Presence of the kanamycin resistance cassette was confirmed by PCR. The *nuoA-N∷kan* and *cyoA-E∷kan* alleles were then moved into strain TR50 by P1 transduction as previously described (Silhavy et al., 1984).

Luminescent transcriptional reporters of *entCEBA* expression were constructed as previously described (Wong et al., 2013). Briefly, the promoter region of the *entCEBA* operon was amplified from E2348/69 or MC4100 using the primers PentCluxF and PentCluxR (table S2) and the high fidelity Phusion DNA polymerase (ThermoFisher) according to the manufacturer’s protocol with the addition of 10% betaine. DNA was separated by electrophoresis on a 1% agarose gel. DNA bands corresponding to the size of the MC4100 *entCEBA* promoter and the E2348/69 *entCEBA* promoter were gel-purified using the GeneJet Gel Purification kit (Fermentas), digested with BamHI and EcoRI (Invitrogen), and ligated upstream of the *luxABCDE* operon in the pJW15 plasmid. PCR and DNA sequencing verified correct insertion of the promoter sequences. To ensure that the reporters reflect accurate expression of the *entCEBA* operon, luminescence was determined under iron-replete and iron-deplete conditions. In accordance with published observations, luminescence was reduced in the presence of iron (data not shown). DNA sequencing was performed by the University of Alberta Molecular Biology Services Unit.

### β-galactosidase assay

For figures 1 and 2, bacteria were grown overnight in LB at 37°C with shaking at 225 rpm. The following day, strains were subcultured at a dilution of 1:100 into fresh LB broth or M9 minimal medium (Difco) and grown for twenty hours at 37°C with shaking at 225 RPM. Where indicated, FeSO_4_ and enterobactin were added to a final concentration of 80μM and 10μM, respectively. As enterobactin is dissolved in 42% DMSO, an equivalent volume of 42% DMSO was added to the control cultures. β-galactosidase activity was measured as previously described (Buelow and Raivio, 2005). Bacteria were pelleted by centrifugation at 2880 × g for 10 minutes. The supernatant was removed, and bacteria were resuspended in 2mL of 1 × Z buffer (10mL of 10x Z buffer [600mM Na_2_HPO_4_•7H_2_O, 400mM NaH_2_PO_4_•H_2_O, 100mM KCl, 10mM MgSO_4_•7H_2_O], 90mL distilled water, 270μL β-mercaptoethanol). 250μL of sample was transferred to a 96 well plate and OD_600_ was measured using the PerkinElmer Wallac Victor^2^ 1420 plate reader. Chloroform and SDS were used to lyse the remaining cells. 5μL of sample was added to 195μL of 1 × Z buffer in a 96 well plate. 50μL of 10mg/mL *o*-nitrophenyl-β-D-galactopyranoside (ONPG) was added, and hydrolysis of ONPG was measured at an absorbance of 420nm (*A*_420_). *A*_420_ was read 20 times with 45 seconds between each reading.

For figure 4 and S1, bacteria were grown overnight in LB at 37°C with shaking at 225 rpm. Bacteria were pelleted by centrifugation at 2880 × g for 10 minutes, washed once in 1mL phosphate-buffered saline, and resuspended in 2mL phosphate buffered saline. 10μL of washed bacteria were spotted onto M9 minimal medium agar containing 0.4% glucose and grown for 24 hours at 37°C. Bacteria were then scraped off the plate using plastic inoculating loops and resuspended in 2mL 1 × Z buffer. β-galactosidase activity was measured as described above.

### NADH Oxidase Assay

After growth overnight in 5mL LB broth with shaking at 225 rpm, bacteria were diluted by a factor of 1:100 into 5mL M9 minimal medium (Difco) containing 0.4% glucose and grown for 20 hours at 37°C. Bacteria were pelleted by centrifugation at 2880 × g for 10 minutes and the pellet was resuspended in 1mL of cold 50mM 4-morpholineethanesulfonic acid (MES) buffer (pH 6.0). Bacteria were pelleted again by centrifugation at 21,130 × g for 1 minute. The supernatant was removed and the wet weight of the bacteria was determined. Bacteria were resuspended in 1mL cold 50mM MES buffer, pH 6.0 and 25μL of protease inhibitor cocktail (Sigma-Aldrich) was added for every 100mg of wet cell weight. Bacteria were then lysed by sonication. Intact cells were removed by centrifugation at 10,000 × g for 30 minutes at 4°C. 100μL of sample was added to 890μL of pre-warmed 50mM MES buffer, pH 6.0 in a 1mL microrespiration chamber and covered with 150μL of light mineral oil to prevent oxygen from dissolving into the medium. The microrespiration chamber was placed in a 30°C water bath for 5 minutes prior to the addition of β-NADH. 100μM β-NADH was added and oxygen concentration was measured every 30 seconds for 10-15 minutes using an oxygen MicroOptode sensor (Unisense). Oxygen concentration at each time point was standardized to the oxygen concentration just prior to the addition of β-NADH. Oxygen consumption for each sample was measured in technical duplicate. The rate of oxygen consumption (% minute^−1^) was calculated from the linear range of the reaction. The average rate of oxygen consumption of two technical replicates was standardized to the amount of total protein added to the microrespiration chamber. Protein concentration for each sample was determined using the Pierce BCA Protein Assay kit (Thermo Scientific).

### Luminescence assay

Bacteria were grown overnight in 2mL LB at 30°C. Bacteria were then diluted at a factor of 1:100 into 2mL M9 minimal medium (Difco) containing 0.4% glucose, 5.34mM isoleucine, and 6.53mM valine and grown at 30°C with shaking at 225 rpm. 200μL of culture was transferred to a black-walled 96 well microtiter plate 8 hours after subculture and luminescence (in CPS) and OD_600_ were measured as described above. Luminescence (expressed in counts per second [CPS]) and OD_600_ were read from the microtiter plate for each sample using the PerkinElmer Wallac Victor^2^ 1420 plate reader. *entCEBA-lux* activity was calculated by subtracting the CPS and OD_600_ values measured from a blank well containing uncultured LB from the raw CPS and OD_600_ values measured for each sample. The normalized CPS was divided by the normalized OD_600_ to account for differences in growth between samples.

### Statistical analysis

Statistical analysis was performed using Prism version 8.2.1 (GraphPad Software). Activity of transcriptional reporters was compared by two-way analysis of variance followed by Sidak’s multiple comparison test.

## Author Contributions

RLG was involved in the design of the study, the acquisition, analysis, and interpretation of the data, and the writing of the manuscript. EAC, JLW, and KAS were involved in the acquisition and analysis of the data. TLR was involved in the conception and design of the study as well as the writing of the manuscript.

## Acknowledgements

We thank Bernard Lemire for help with designing the NADH oxidase assay. We are thankful to the members of the Raivio lab for their comments. This work was funded by operating grant MOP 342982 awarded to T.L.R from the Canadian Institutes of Health Research.

