## Supplementary Material for "Impaired Efflux of the Siderophore Enterobactin Induces Envelope Stress in *Escherichia coli*"

### Materials and Methods

#### Sequence alignment

The DNA sequence of the *entCEBA* promoter region in enteropathogenic *E. coli* strain E2348/69 and *E. coli* K-12 strain MC4100 obtained from the cloned pJW15-P*entCEBA*$_{EPEC}$ and pJW15-P*entCEBA*$_{K-12}$ reporter plasmids. DNA sequencing was performed by the Molecular Biology Services Unit at the University of Alberta. The DNA sequence of the insert for the pJW15-P*entCEBA*$_{EPEC}$ and pJW15-P*entCEBA*$_{K-12}$ was compared to the published genome of MG1655 and E2348/69, respectively, and was found to be 100% identical (data not shown). DNA sequences were aligned using Multialin (http://multalin.toulouse.inra.fr/multalin).

### Figures and Tables

#### Table S1. Bacterial strains and plasmids used in this study

| Strain or plasmid | Description | Source or reference |
|---|---|---|
| *Strains* | | |
| MC4100 | F’ *traD36 lacI$^q$ Δ(lacZ)M15 proA$^+$B$^+$/* e14 (McrA$^-$) *Δ(lac-proAB) thi gyrA96* (Nal$^r$) *endA1 hsdR17*(r$_k^-$ m$_k^+$) *relA1 supE44 recA1*; Str$^R$ | (Casadaban, 1976) |
| TR10 | MC4100 *cpxA24*; Amk$^R$ | (Raivio *et al.*, 1999) |
| TR50 | MC4100 λRS88[*cpxP’-lacZ$^+$*]; Str$^R$ | (Raivio and Silhavy, 1997) |
| TR51 | MC4100 *cpxR::spc*; Spc$^R$ | (Raivio *et al.*, 1999) |
| JW0585 | BW25113 Δ*entC::kan*; Kan$^R$ | (Baba *et al.*, 2006) |
| JW1095 | BW25113 Δ*ndh::kan*; Kan$^R$ | (Baba *et al.*, 2006) |
| JW5503 | BW25113 Δ*tolC::kan*; Kan$^R$ | (Baba *et al.*, 2006) |
| DY378 | W3110 λcI857 Δ(*cro-bioA*) | (Yu *et al.*, 2000) |
| E2348/69 | Prototypical EPEC O127:H6 strain; Str$^R$ | (Levine *et al.*, 1978) |
| ALN195 | E2348/69 *cpxA24*; Str$^R$ Amk$^R$ | (MacRitchie *et al.*, 2008) |
| RG222 | E2348/69 Δ*cpxRA*; Str$^R$ | (Guest *et al.*, 2017) |
| EC3 | TR50 Δ*tolC*; Str$^R$ | This study |
| EC4 | TR50 Δ*tolC* Δ*entC::kan*; Kan$^R$ | This study |
| RG244 | TR50 Δ*entC::kan*; Kan$^R$ | This study |
| RG249 | TR50 Δ*ndh::kan*; Kan$^R$ | This study |
| RG250 | TR50 Δ*tolC* Δ*ndh::kan*; Kan$^R$ | This study |
| RG280 | TR50 Δ*cpxA::kan*; Kan$^R$ | This study |
| RG281 | TR50 Δ*tolC* Δ*cpxA::kan*; Kan$^R$ | This study |
| RG383 | DY378 Δ*nuoABCDEFGHIJKLMN::kan*; Kan$^R$ | This study |
| RG392 | TR50 Δ*nuoABCDEFGHIJKLMN::kan*; Kan$^R$ | This study |
| RG397 | DY378 Δ*cyoABCDE::kan*; Kan$^R$ | This study |

| | | |
|---|---|---|
| RG436 | TR50 Δ*cyoABCDE::kan*; $Kan^R$ | This study |
| RG479 | TR50 Δ*tolC* Δ*nuoABCDEFGHIJKLMN::kan*; $Kan^R$ | This study |
| RG480 | TR50 Δ*tolC* Δ*cyoABCDE::kan*; $Kan^R$ | This study |
| *Plasmids* | | |
| pFLP2 | Broad host-range plasmid expressing the FLP recombinase from a temperature sensitive promoter; $Amp^R$ | (Hoang *et al.*, 1998) |
| pJW15-P*entCEBA*$_{K\text{-}12}$ | pJW15 luminescence reporter plasmid containing the MC4100 *entCEBA* promoter; $Kan^R$ | This study |
| pJW15-P*entCEBA*$_{\text{EPEC}}$ | pJW15 luminescence reporter plasmid containing the E2348/69 *entCEBA* promoter; $Kan^R$ | This study |

**Table S2. Oligonucleotide primers used in this study**

| Primer name | Sequence* |
|---|---|
| PentFEcoRI | 5'-TTTTGAATTCCTGAACTGCGGCTATTCCTG-3' |
| PentRBamHI | 5'-TTTTGGATCCTACTTCCTCAGCCAGTGACG-3' |
| K12-cyoKOF | 5'-CCACACACTTTAAACGCCACCAGATCCCGTGGAATTGAGG TCGTTAAATGATTCCGGGGATCCGTC-3' |
| K12-cyoKOR | 5'-CGTAGCACCTTTTTAATAGAGAGGTTTTGTTACCACACAGCA GCCAGCAGTGTAGGCTGGAGCTGC-3' |
| K12nuoKOF | 5'-CTGCCGTGAAGAGCAGTGAATCTGGCGCTACTTTTGATGAGT AAGCAATGATTCCGGGGATCCGTC-3' |
| K12nuoKOR | 5'-GGCGGCTTTCTGACTTACAAAGTAACAGATTACATCAGCGGC ATTGCCAATGTAGGCTGGAGCTGC-3' |

*Underlining denotes a restriction enzyme sequence (BamHI: GGATCC; EcoRI: GAATTC)

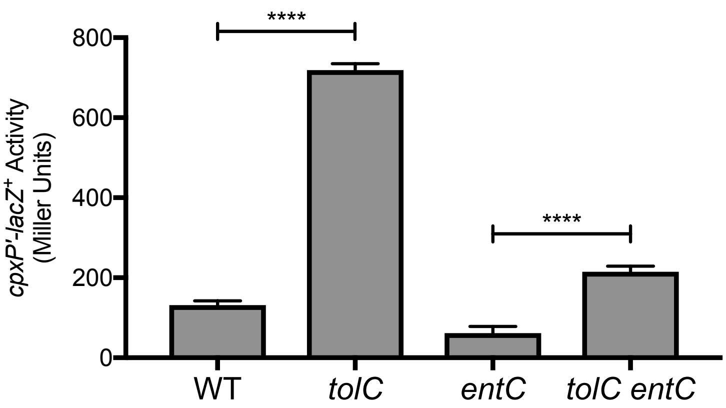

**Figure S1: Enterobactin accumulation activates the Cpx response in the *tolC* mutant.** After growth overnight in LB broth, bacteria were washed and resuspended in phosphate buffered saline. 10µL of culture was spotted onto M9 minimal medium agar containing 0.4% glucose and grown at 37°C for 24 hours. Bacteria were scraped off the agar surface using inoculating loops and resuspended in 1 x Z buffer. *cpxP-lacZ* activity was measured as described in the materials and methods section of the main text. Data represent the means and standard deviations of three biological replicates. Asterisks indicate a statistically significant difference in *cpxP-lacZ* activity between the indicated strains (****, $P \leq 0.0001$ [two-way ANOVA with Sidak's post-hoc test]).

**A.**

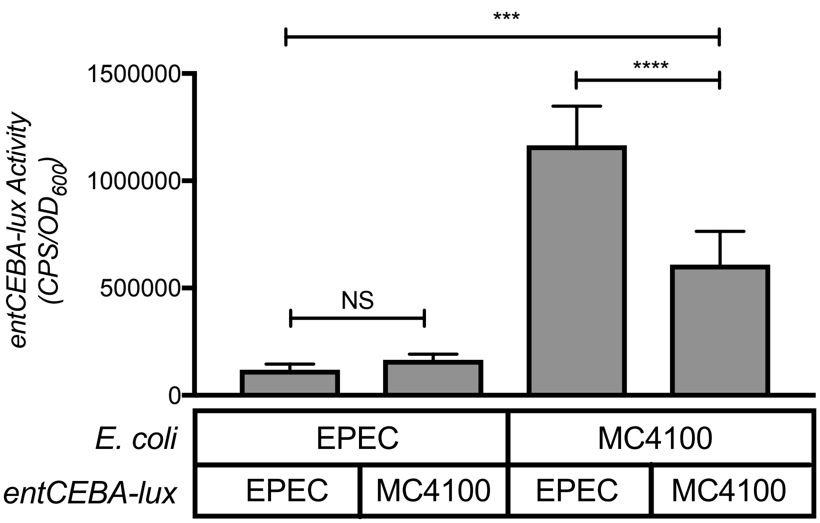

**B.**

```
  MC4100  CCTGAACTGCGGCTATTCCTGAAAGCAAAAGTCCTGTTAATAGAAGGGCG
E2348/69  CCTGCACTGCGGCTATTCCTGAAAGCAAAAGTCCTGTTAATAGAAGGGCG

  MC4100  TTGCGGTAGAGCGGGGCGAGTCTCACAAATCAGCTTCCTGTTATTAATAA
E2348/69  GTGCGGTAGAGCGGGGCGAGTCTCACAAATTAGCTTCCTGTTATTAATAG

  MC4100  GGTTAAGGGCGTAATGACAAATTCGACAAAGCGCACAATCCGTCCCCTCG
E2348/69  AGTTAA--------------------------------------------

  MC4100  CCCCTTTGGGGAGAGGGTTAGGGTGAGGGGAACAGCCAGCACTGGTGCGA
E2348/69  --------------------------------------------------

  MC4100  ACATTAACCCTCACCCCAGCCCTCACCCTGGAAGGGAGAGGGGGCAGAAC
E2348/69  --------------------------------------------------

  MC4100  GGCGCAGGACATCACATTGCGCTTATGCGAATCCATCAATAATGCTTCTC
E2348/69  ------------------------------------------TGCTTCTC

  MC4100  ATTTTCATTGTAACCACAACCAGATGCAACCCCGAGTTGCAGATTGCGTT
E2348/69  ATTTTCATTGTAACCACAAACAGATGCAACCCCGAGTTGCAGATTGCGTT

  MC4100  ACCTCAAGAGTTGACATAGTGCGCGTTTGCTTTTAGGTTAGCGACCGAAA
E2348/69  ACCTCAAGAGTTGACATAGTGCGCGTTTGCTTTTAGGTTAGCGACCGAAA

  MC4100  ATATAAATGATAATCATTATTAAAGCCTTTATCATTTTGTGGAGGATGAT
E2348/69  ATATAAATGATAATCATTATTAAAGCCTTTATCATTTTGTGGAGGATGAT

  MC4100  ATGGATACGTCACTGGCTGAGGAAGTA
E2348/69  ATGGATACGTCACTGGCTGAGGAAGTA
```

**Figure S2: Comparison of *entCEBA* expression in MC4100 and EPEC.** (A) Activity of the enteropathogenic *E. coli* (EPEC) *entCEBA-lux* reporter and the MC400 *entCEBA-lux* reporter in wildtype EPEC or wildtype MC4100. Bacteria were grown overnight in LB broth. The following day, bacteria were subcultured into M9 minimal medium containing 0.4% glucose, 5.34mM isoleucine, and 6.53mM valine at a dilution factor of 1:100. *entCEBA-lux* expression was measured after 8 hours of growth at 30$^{o}$C as described in the materials and methods section of the main text. Data represent the means and standard deviations of five biological replicates. Asterisks indicate a statistically significant difference between the indicated strains (****, $P \leq 0.0001$; ***, $P \leq 0.001$ [one-way ANOVA with Sidak's post-hoc test]). NS indicates no statistically significant difference in *entCEBA-lux* reporter activity. (B) Alignment of the *entCEBA* promoter region between MC4100 and E2348/69. DNA sequence of the *entCEBA* promoter DNA was determined by sequencing the insert of the pJW15-P*entCEBA*$_{K\text{-}12}$ and pJW15-P*entCEBA*$_{EPEC}$ plasmids. The DNA sequences were aligned using Multialin (http://multalin.toulouse.inra.fr/multalin). Bolded sequences represent the -35 box (TTGACA), the -10 box (TAGGTT), and the start codon (ATG), which were identified using the Ecocyc database (http://ecocyc.org) (Keseler *et al.*, 2011). Red sequences denote single base pair changes; -, absence of a base pair.
